# Loss of the Long Non-coding RNA OIP5-AS1 Exacerbates Heart Failure in a Sex-Specific Manner

**DOI:** 10.1101/2021.01.28.428540

**Authors:** Aowen Zhuang, Anna C. Calkin, Shannen Lau, Helen Kiriazis, Daniel G. Donner, Yingying Liu, Simon T. Bond, Sarah C. Moody, Eleanor A.M. Gould, Timothy D. Colgan, Sergio Ruiz Carmona, Michael Inouye, Thomas Q. de Aguiar Vallim, Elizabeth J. Tarling, Gregory A. Quaife-Ryan, James E. Hudson, Enzo R. Porrello, Paul Gregorevic, Xiao-Ming Gao, Xiao-Jun Du, Julie R. McMullen, Brian G. Drew

## Abstract

**Background:** Long ncRNAs (lncRNAs) are known to influence numerous biological processes including cellular differentiation and tissue development. They are also implicated in the maintenance, health and physiological function of many tissues including the heart. Indeed, manipulating the expression of specific lncRNAs has been shown to improve pathological cardiac phenotypes such as heart failure. One lncRNA studied in various settings is OIP5-AS1 (also known as *1700020I14Rik* and *Cyrano*), however its role in cardiac pathologies remains mostly uncharacterised.

**Methods:** We used data generated from FACS sorted murine cardiomyocytes, human iPSC derived cardiomyocytes, as well as heart tissue from various animal models to investigate OIP5-AS1 expression in health and disease. Using CRISPR we engineered a global OIP5-AS1 knock out (KO) mouse model and performed cardiac pressure overload experiments to study heart failure in these animals. RNA-sequencing of left ventricles provided mechanistic insight between WT and KO mice.

**Results:** We demonstrate that OIP5-AS1 expression is regulated during cardiac development and cardiac specific pathologies in both rodent and human models. Moreover, we demonstrate that global female OIP5-AS1 KO mice develop exacerbated heart failure, but male mice do not. Transcriptomics and gene set enrichment analysis suggests that OIP5-AS1 may regulate pathways that impact mitochondrial function.

**Conclusions:** OIP5-AS1 is regulated in cardiac tissue and its deletion leads to worsening heart function under pressure overload in female mice. This may be due to impairments in mitochondrial function, highlighting OIP5-AS1 as a gene of interest in sex-specific differences in heart failure.

## Introduction

Since the advent of deep-sequencing and genome mapping, it has become clear that genetic control of cellular function is evidently more complex than the classical model of DNA➔RNA➔protein. This dogma was initially re-examined upon the discovery of miRNAs, which were one of the first classes of non-coding (nc)RNAs demonstrated to regulate cellular disease pathways. Since then, the ncRNA field has expanded rapidly with the subsequent discovery of a wide range of ncRNA classes, including potentially thousands of predicted long non-coding RNAs (lncRNAs) [1]. LncRNAs are those described to be >200 nucleotides long; however, most are several kilobases in length and are topographically similar in many ways to protein coding mRNAs [2]. *Bona fide* lncRNAs do not code for functional proteins and are therefore proposed to have a vast array of functions including acting as transcriptional regulators, structural scaffolds or RNA sponges [3,4]. Although lncRNAs were discovered more recently, exciting studies have already emerged to suggest that they are likely promising targets for therapeutic and biomarker applications. Indeed, lncRNAs are often more cell-type specific and tightly regulated than protein coding RNAs (mRNA) [5].

Of particular interest to the current study, are several lncRNAs that have been linked with pathological conditions in cardiac muscle [6]. The molecular regulation of cardiac commitment and development has been an intense area of research for many years. The discovery of lncRNAs has thus pioneered a new area of investigation in cardiac biology, with several groups identifying cardiac specific lncRNAs that are involved in almost every facet of cardiac commitment, development and function. The first detailed mechanistic description of a cardiac specific lncRNA was Braveheart (*bvht*) [7], which was shown to be important in cardiac lineage commitment through its actions on MesP1, the master regulator of cardiovascular lineage commitment. Since then, many lncRNAs have been implicated in cardiac commitment and differentiation including *Fendrr*, *SRA* and *Novlnc6*, all of which affect the activity of lineage specific transcriptional pathways [8–10]. Furthermore, the expression of some lncRNAs such as *MIAT, LIPCAR, Mhrt* and *CHRF* are also strongly associated with cardiovascular disease [11–14]. Specifically, MyHeart (*Mhrt*) and *CHRF* exhibit significantly altered expression in the setting of cardiac hypertrophy [11,14]. Importantly, mice with reconstitution of *Mhrt* expression in the setting of pathological hypertrophy, displayed less cardiac dysfunction than wild type mice, providing evidence that lncRNAs have promising therapeutic potential in the heart [11].

Here, we investigate a lncRNA known as OIP5-AS1 (also known as *1700020I14Rik* and Cyrano), which has been reasonably well studied in the brain and ES cells, but there is limited data on its roles in cardiac pathologies such as heart failure. We reveal that OIP5-AS1 expression is enriched in striated muscle and differentiating cardiomyocytes, and its expression is reduced in the setting of heart failure. Furthermore, we show that global loss of OIP5-AS1 in mice leads to poorer outcomes following pressure-overload induced heart failure, but only in female mice. Thus, our studies demonstrate a previously unrecognised sex-specific function of OIP5-AS1 in the heart.

## Methods

### Animals

All animal experiments were approved by the Alfred Research Alliance (ARA) Animal Ethics Committee (E/1769/2017/B), and performed in accordance with the NH&MRC of Australia guidelines for the care and use of laboratory animals. OIP5-AS1 knockout (KO) mice were generated by the Australian Phenomics Network with deletion being achieved by engaging CRISPR/Cas9 technology in single cell embryos, using gRNAs targeting either end of the genomic sequence of the *1700020I14Rik* (OIP5-AS1) gene. Edited C57BL/6J embryos were implanted into pseudo-pregnant C57BL/6J female mice, and founder offspring were sequenced to confirm genetic modification. Founders were subsequently mated to confirm germline editing and bred for >5 generations to C57BL/6J mice that had not been genetically manipulated. Cohorts for experimental studies were bred and sourced through the ARA Animal Centre and randomly allocated to groups. After weaning at 4 weeks of age, wild type and KO male and female mice were matched for sex and body weight. Because male mice are commonly bigger than female mice, male mice were generally 1-2 weeks younger than female at time of surgery, in order to ensure that surgeries were performed on weight-matched animals. All animals were housed at 22°C on a 12hr light/dark cycle with *ad libitum* access to food (standard rat and mouse chow, Specialty feeds, Australia) and water, with cages changed weekly.

### Transverse-Aortic Constriction (TAC)

Transverse restriction of aortic outflow was performed on 7-10 week old animals as previously described [15,16]. Briefly, animals were anesthetized with a mixture of ketamine, xylazine, and atropine (10, 2, and 0.12 mg/100 g, respectively, ip), intubated via the oral cavity, and ventilated. Following a sternotomy, the transverse aorta between the right innominate and left carotid arteries was dissected and banded with a 26-gauge blunt needle using a 5-0 silk suture. The probe was then removed allowing for the suture to remain in place and thus permit chronic constriction of the aorta. With the use of this diameter needle, aortic diameter was predicted to be reduced by 50–55%, which leads to an approximately 70% reduction in cross-sectional area. This procedure was performed similarly on both male and female mice between the ages of 7-10 weeks, which allowed us to weight match the animals and thus assume a mouse of a similar weight would have a similar sized aorta and percent constriction. Previously, other studies have used mice of the same age when comparing sex differences, meaning that the bigger male mice would likely receive a more severe aortic restriction in age-matched animals, resulting in more severe disease. Sham-operated mice underwent the same surgical procedure as TAC treated mice, but no suture was place around the aorta to restrict flow.

### Intra-aortic and LV pressure analysis

Blood pressure and intra-cardiac pressure was assessed by a catheter placed into the right carotid artery (proximal to the stenotic site) and advanced into the LV as previously described [17]. Mice were anesthetized using isoflurane at 2-4% and placed in the supine position on a heating pad, and the right main carotid artery was dissected. A micro-tipped transducer catheter (1.4F, Millar Instrument Co) was inserted into the artery and measurements including aortic blood pressure and LV pressures were recorded.

### Echocardiography

Echocardiography was performed on mice anaesthetised with isoflurane (1.75%) at baseline and at the end of the 8-week study using a 15-MHz linear transducer L15-7io with a Philips iE33 Ultrasound Machine (North Ryde, NSW, Australia). Data were analysed and verified by two independent researchers according to QC procedures and validation measures as outlined previously {Donner, 2018 #1251}.

### Quantitative PCR (qPCR)

RNA for qPCR analysis was isolated from tissues as previously described [19,20]. Briefly, tissues were homogenised in RNAzol reagent and precipitated using isopropanol. cDNA was generated from 1μg of RNA using MMLV reverse transcriptase (Invitrogen) according to the manufacturer’s instructions. qPCR was performed on 10ng of cDNA using the SYBR-green method on a QuantStudio 7 Flex (ThermoFisher Scientific) using gene specific primers. Quantification of a given gene was expressed by the relative mRNA level compared with control, which was calculated after normalisation to the housekeeping gene 36B4 (*Rplp0*) using the delta-CT method. Primers were designed to span exon-exon junctions where possible and were tested for specificity using BLAST (Basic Local Alignment Search Tool; National Centre for Biotechnology Information) (see **Table 1** for Primer Details). In order to account for the two main variants of OIP5-AS1 expressed in most cell types, including striated muscles, we designed 4 sets of primers. Two of these primer sets recognized all variants, whilst another two recognized only the most abundant, and longest variant of OIP5-AS1 (known as Oip5os1-202 in *mus musculus*). Amplification of a single amplicon was estimated from melt curve analysis, ensuring only a single peak and an expected temperature dissociation profile were observed.

**Table 1:**
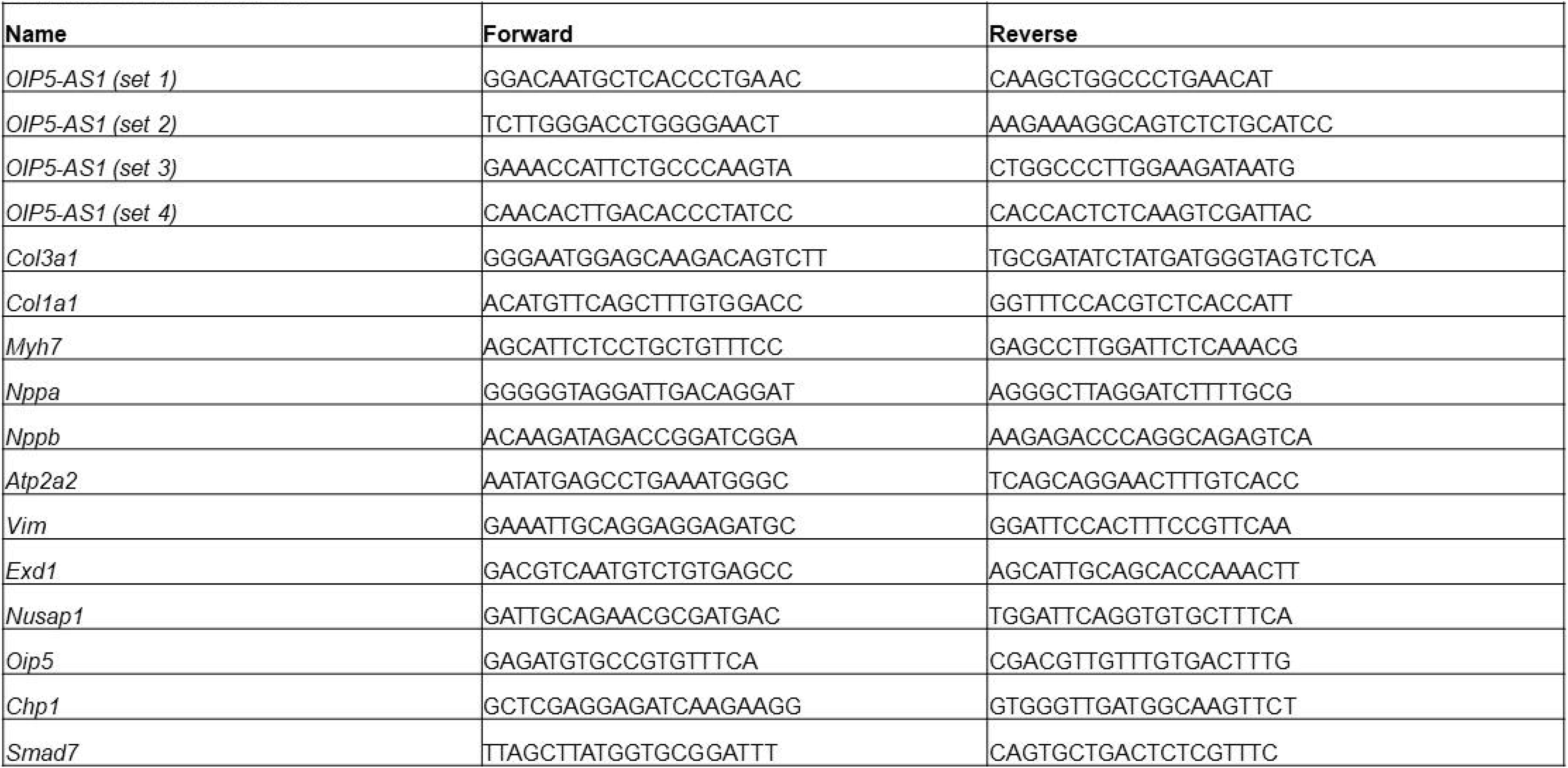
qPCR Primer Sequences. Nucleotide sequence (5’ – 3’) of the primers used for quantitative PCR analysis using the SYBR green method.

### RNA-sequencing Analysis

RNA was isolated from LV tissue using RNAzol reagent and purified using columns according to the manufacturer’s instructions (Zymo Research). RNA integrity was evaluated using the Agilent Tape Station 2200 according to the manufacturer’s instructions (Agilent). RNA libraries were prepared using Kapa Stranded RNA-seq kits on samples with a RIN>0.8 according to the manufacturer’s instructions (Roche). Library quantities were determined using QUBIT (ThermoFisher) and equal amounts of all 24 libraries were pooled and run across 2 lanes using an Illumina HiSeq 2500 Sequencer. FASTQ sequencing data were demultiplexed and aligned to the *Mus Musculus* mm10 genome using STAR Software (V2.7.1) with default parameters. Mapped reads were counted using *Featurecounts* against an mm10 reference file. QC and batch effect analyses were performed using edgeR and DEseq2 bioconductor packages in R.

### Pathway and Network Analysis

Enrichment analysis and gene ontology was performed using the Database for Annotation, Visualization and Integrated Discovery (DAVID v.6.8) hosted by the National Institute of Allergy and Infectious Diseases (NIAID), NIH, USA [21]. Cluster analysis from human iPSC-derived cardiomyocyte RNA-sequencing were derived by analysing gene sets correlated with OIP5-AS1 expression. Cluster analysis from RNA-sequencing data were derived from genes sets significantly altered between WT and KO animals. These datasets were separately analysed using GSEA (v3.0) [22,23] and functional enrichment analysis was mapped using g:Profiler (v95_eg42_p13_f6e58b9) with g:SCS multiple testing correction method applied with a significance threshold of p<0.05 [24,25]. Enrichment and cluster analysis were mapped to a network of the curated MSigDB C5 gene set collection (nodes)[23].

### Histology

LV tissues were fixed in paraformaldehyde overnight before being placed in 70% ethanol. Tissues were subsequently embedded in paraffin blocks and 5μm sections were cut and mounted on glass slides. After dewaxing and hydration, sections were stained using picrosirius red (Sigma) to visualise cell collagen and fibrosis abundance. Slides images were captured using Olympus Slide scanner VS120 (Olympus, Japan) and viewed in the supplied program (OlyVIA Build 13771, Olympus, Japan). Whole tissue slides were quantified based on threshold analysis in Fiji [26].

### Primary Endpoints and Data Inclusion/Exclusion Criteria

Our primary endpoint was to test if OIP5-AS1 KO affected heart function in either male or female mice, compared to their relative WT controls. For *in vitro* and basal animal phenotyping data, individual data points were excluded if they were technically implausible or a methodological error had resulted in a spurious outcome. Analyses from animals following TAC surgery were excluded if animals were found dead from acute heart failure overnight (samples compromised), did not recover from surgery (2 female KO mice post-TAC) or technical/analytical problems were identified (compromised RNA, failed analysis, improper tissue collection, equipment failure). For echocardiography, data was excluded if heart rates were outside of 450-650bpm or animals were too ill to undergo anaesthesia. Echocardiography outputs were only included in final analysis if we had a full pre- and post-surgery dataset, meaning that numbers were lower in groups where TAC surgery induced a more severe heart failure (i.e. female KO mice).

### Statistical Analysis and Sample Size

Our primary endpoint was to identify if there were differences between WT and KO mice within each sex (and not between sexes), therefore sample size was determined accordingly. All animal and laboratory data underwent blinding and randomization at time of collection and during technical analysis. Data were expressed as mean ± standard error of the mean (SEM), unless otherwise stated. All statistical analyses of animal and laboratory based experiments were performed using PRISM7 software. Normally distributed data in cell culture experiments were compared by paired students t-test whilst animal studies were analysed by one-way and two-way ANOVA with testing for multiple comparisons between WT and KO animals of the same sex. As our primary endpoint was not to determine if male and females were statistically different from each other, we did not perform multiple comparison testing with all four groups. In these analyses, a p-value of p<0.05 was considered statistically significant.

## Results

### OIP5-AS1 is enriched in striated muscles

OIP5-AS1 is located between the two protein coding genes *Chp1* and *Oip5* on chromosome 2 (**Supplemental Figure 1A**), and is a *bona fide* lncRNA based on its absence in ribosomal transcriptome profiling [27]. It is called OIP5-AS1 because of its proximity and opposing topographical locality to the *OIP5* gene, which is similar to that observed in the mouse [28]. OIP5-AS1 has been the topic of a number of studies in lab-based systems such as zebrafish and embryonic stem cells (ES cells). Its function in these systems have been described to modulate the abundance of specific miRNAs (miRs) to influence pathways important in proliferation [29], self-renewal [30] and differentiation [31]. However, functional genetic studies have failed to demonstrate a link between loss of OIP5-AS1 expression and robust phenotypes in animal systems, other than mild malformation of the neural tube and nasal placodes in zebrafish embryos – hence its alternative name, *Cyrano* [27]. Moreover, almost none of these phenotypes have translated across vertebrate species, with mild to no phenotype identified in knock-down or partial KO models in higher order mammals (i.e. mice)[28].

Using Genomic Evolutionary Rate Profiling (GERP) data, we and others [27,28] demonstrate that unlike the majority of lncRNAs, OIP5-AS1 harbours several regions of high homology in its nucleotide sequence between mammalian species including humans. We also confirm previous findings that a small region of exon 3 is 100% conserved across vertebrates (**Supplemental Figure 1B**; blue shaded area). This is uncommon for lncRNAs, which often exhibit poor sequence conservation between species [32].

In the current study we were interested to know if OIP5-AS1 has functional relevance in the heart, especially given that deposited gene expression data from NCBI (**Figure 1A**) [33] and transcriptomics data from our lab (**Supplementary Table 1**)[34] and others [28], demonstrate an enrichment for OIP5-AS1 in striated muscles (diaphragm, muscle and heart), as well as in the brain. Indeed, our transcriptomics data (from skeletal muscle) demonstrated that OIP5-AS1 is expressed within the top 500 most abundant genes (out of ~7000 detected), together with several other previously annotated lncRNAs such as *Malat1*, *H19* and *Rhit1 (Nctc1)* [35,36].

**Figure 1:**
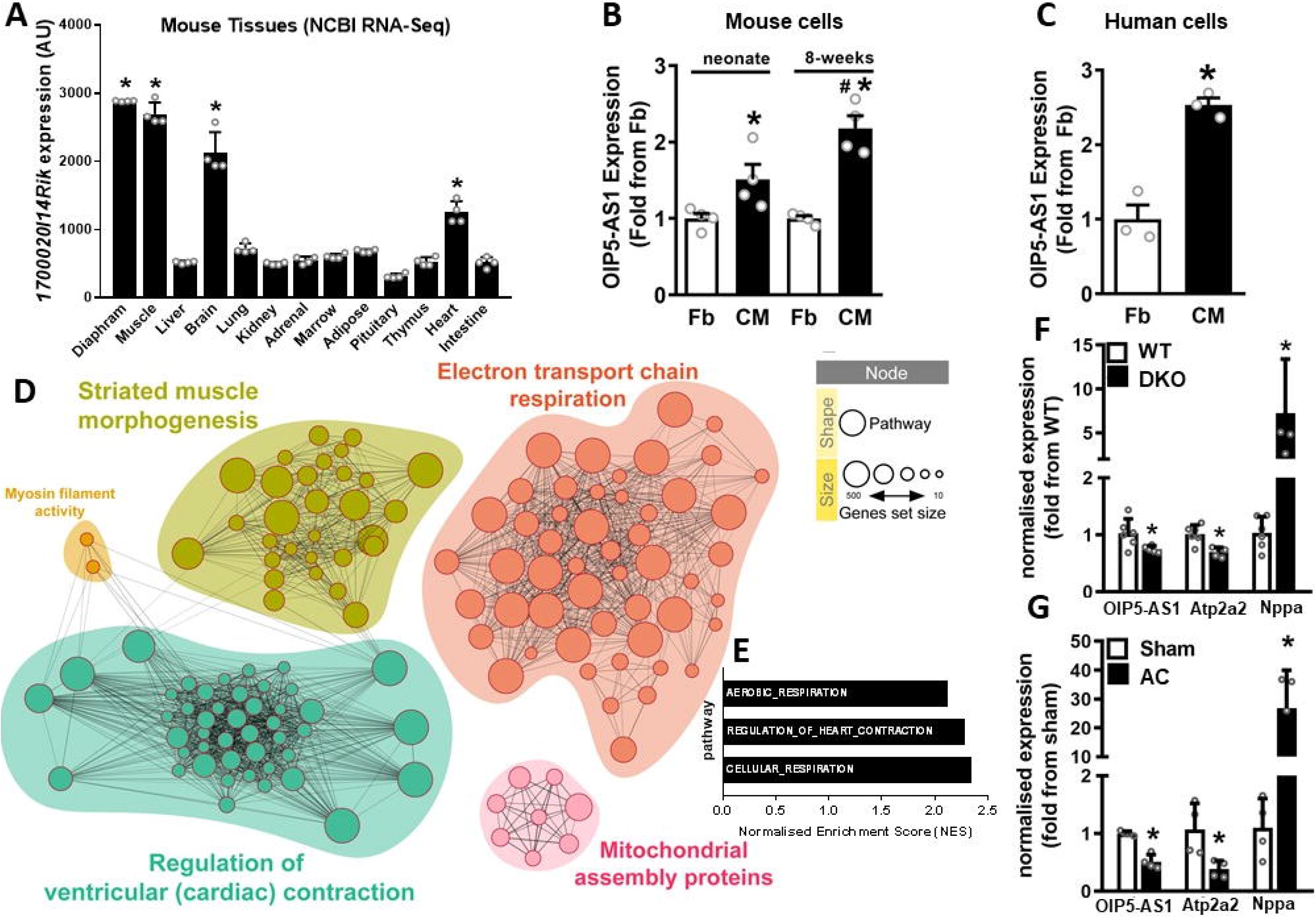
OIP5-AS1 is enriched in developing cardiac muscle, and is altered during cardiac disease. **A**. Expression of OIP5-AS1 across mouse tissues sourced from NCBI Gene Expression Omnibus (GEO, GSE24207), n=4/group, mean±SEM, * p<0.001 from liver expression. **B**. Expression of OIP5-AS1 as determined by RNA-sequencing in fibroblasts (Fb) and cardiomyocytes (CM) digested from mouse hearts of neonate and 8-week old mice (n=4/group, mean±SEM, * p<0.05 versus neonate Fb; # p<0.05 versus neonate CM) from GEO deposited dataset GSE95764, and in **C**. Fb and CM differentiated from human iPSCs (n=3/group, mean±SEM, * p<0.05 versus Fb). **D**. Network analysis of genes sets correlated with OIP5-AS1 expression as determined by RNA-sequencing in mouse cardiomyocytes versus fibroblasts from datasets in panel C. Enriched networks include cardiac contraction, muscle morphogenesis and mitochondrial pathways (assembly and electron transport chain (ETC)). For a description of each code shown within the nodes, see **Supplementary Table S2**. **E**. Bar graph depicting most highly enriched pathways associated with OIP5-AS1 expression in mouse cardiomyocytes. **F**. qPCR determined expression of OIP5-AS1, *Atp2a2* (SERCA2) and *Nppa* (ANP) in hearts from WT or utrophin/dystrophin double KO (DKO) mice (n=8/group, mean±SEM, * p<0.05 versus WT) [40], and **G**. mice that have undergone sham or aortic constriction (AC); n=4-6/group, mean±SEM, * p<0.05 versus sham). For A, B, C, F & G, a non-parametric one-way ANOVA with multiple comparisons correction (Dunnet’s) was used.

### OIP5-AS1 is regulated during development of cardiac muscle

To determine whether OIP5-AS1 expression is regulated in cardiac muscle, we examined its expression in various models and tissues. Firstly, we sought to investigate whether its expression was enriched in cardiomyocytes (CM) relative to other cells types in the heart, and therefore investigated its expression profile in fibroblasts (Fb) and cardiomyocytes (CM) isolated from the murine heart. To do this, we examined RNA-seq data (GEO dataset GSE95764) from mouse Fbs and CMs that were FACS sorted from digested neonatal and adolescent mouse hearts (**Figure 1B**)[37]. These data demonstrated that OIP5-AS1 expression was higher in CMs of both neonates and adolescents, with an increasing expression observed in the adolescent (8-weeks) heart. We also investigated the expression of OIP5-AS1 in an *in vitro* human model of cardiomyogenesis. RNA-seq data from human fibroblasts and cardiomyocytes induced from the same pluripotent stem cell (iPSCs) population, again demonstrated an increased expression of OIP5-AS1 in CM compared to fibroblasts (**Figure 1C**). Collectively, these findings provide evidence that OIP5-AS1 expression is upregulated in cardiomyocytes compared to fibroblasts in both rodent and human cells

Given that OIP5-AS1 appears to be regulated during cardiomyogenesis, we performed computational analyses in an attempt to predict the potential role that OIP5-AS1 might play in cardiomyocytes. Using RNA-seq data from the mouse fibroblasts and CM datasets described above, we demonstrated that OIP5-AS1 transcript associated with gene networks representative of ventricular contraction, muscle morphogenesis and mitochondrial function (**Figure 1D**, for description of enriched GO terms see **Supplementary Table 2**). Enrichment analysis reveals a link to pathways that are consistent with alterations in respiration and heart contraction (**Figure 1E**), providing evidence of a link between OIP5-AS1 and cellular pathways involved in heart energetics.

### OIP5-AS1 expression is regulated in the setting of disease

Given the results above, we considered whether OIP5-AS1 might also be altered in cardiac disease – a setting where developmental processes and cardiac energetics are often dysregulated. Alterations in the expression of lncRNAs have been observed in several models of cardiovascular disease such as cardiomyopathy and myocardial infarction [6]. Indeed, OIP5-AS1 expression was previously reported to be reduced in the hearts of rats that had suffered from a myocardial infarction[38]. To demonstrate if OIP5-AS1 expression was specifically affected in heart failure, we chose to analyse tissues from two existing murine heart failure models. This included the mild model of heart failure, the utrophin/dystrophin double knock-out (DKO) mouse[39],[40]. Analysis of hearts from these mice demonstrated a reduction in SERCA2 (*Atp2a2*) and an increase in ANP (*Nppa*) mRNA expression compared to wild type mice, consistent with molecular signatures of heart failure (**Figure 1F**). We also observed a reduced expression of OIP5-AS1, suggesting that OIP5-AS1 is downregulated in the setting of mild heart failure. Next, we studied hearts from mice that had undergone aortic constriction (AC) induced by cardiac pressure overload [41,42], which is a severe model of heart failure. In this model, we observed robust changes in the cardiac expression of SERCA2 and ANP, as well as a consistent reduction in OIP5-AS1 expression (**Figure 1G**). Collectively, these data provide evidence that OIP5-AS1 expression is downregulated in the setting of heart failure and cardiomyopathy in mice.

### Generation of an OIP5-AS1 knockout mouse

The above findings are mostly associative, and thus do not specifically demonstrate a direct role for OIP5-AS1 in cardiac pathologies. Therefore, to investigate whether OIP5-AS1 is causally linked to cardiac dysfunction and disease, we generated an OIP5-AS1 global knockout mouse using CRISPR/Cas9 technology in C57BL/6J mice by deleting the OIP5-AS1 gene (**Figure 2A**). These mice were viable and subsequently bred with wild-type (WT) C57BL/6J mice for 5 generations before generating cohorts of WT and KO mice. Using qPCR analysis, we demonstrated complete ablation of OIP5-AS1 expression in all muscle tissues tested in homozygous null mice (KO) (**Figure 2B**), confirming successful generation of the model. Young adult (~10 weeks of age) KO mice were phenotypically unremarkable and displayed no differences in body weight or organ weights compared to WT mice (**Figure 2C**). These findings are consistent with that from Kleaveland and colleagues who generated a partial exon 3 OIP5-AS1 KO mouse that also displayed no overt basal phenotype [28]. Furthermore, basal phenotyping of heart function in our model using echocardiography, demonstrated no difference in left ventricle dimensions or function between WT and KO male or female mice at 10 weeks of age (**Table 2**).

**Figure 2:**
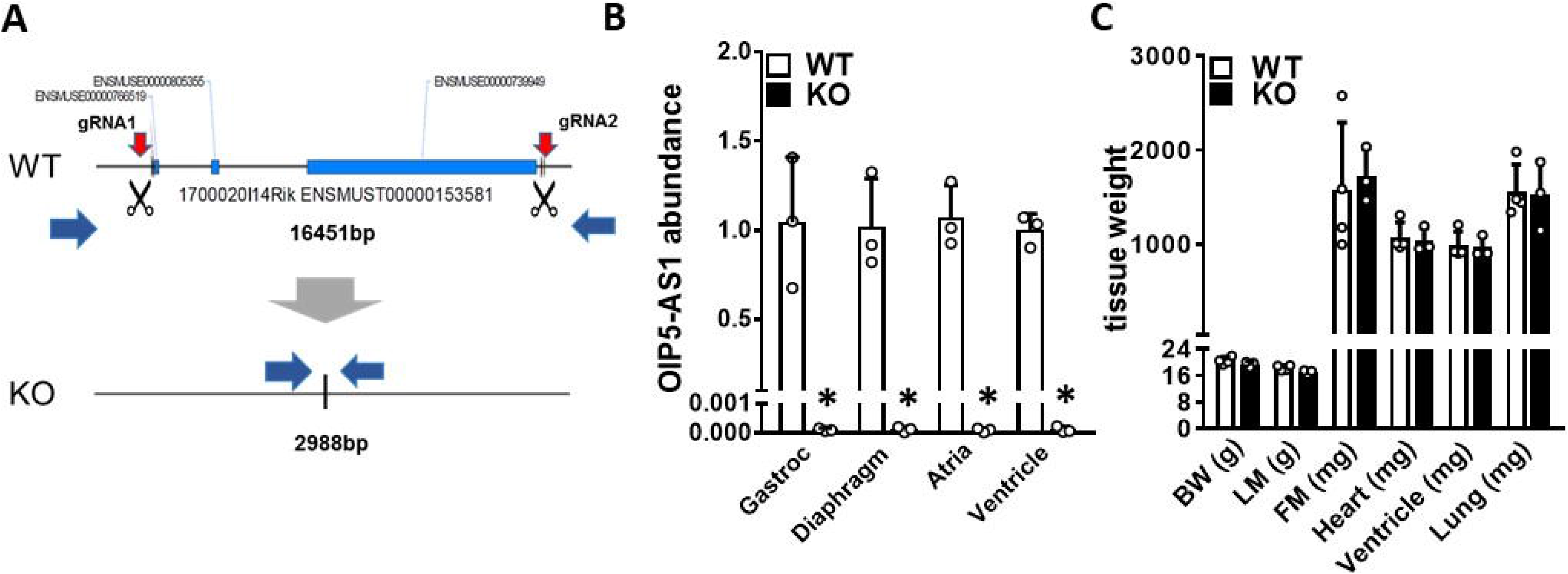
Generation and Basal Phenotyping of an OIP5-AS1 KO mouse model. **A**. Schematic outlining the CRISPR/Cas9 approach employed to delete the OIP5-AS1 gene (~13.4kb) from the C57BL/6J genome. Two guide RNAs (gRNA1&2) were designed to bind at either end of the gene locus (red arrows) after which Cas9 induced removal of the entire gene by non-homologous repair mechanisms. **B**. qPCR analysis of OIP5-AS1 gene expression from four different muscle tissues from 10-week old male WT and OIP5-AS1 KO (KO) mice (n=5/group, mean±SEM, * p<0.05 versus WT). Gastroc = *Gastrocnemius* **C**. Body mass and organ weights of 10-week old female WT and KO mice (n=3-5/group, mean±SEM). BW = body weight, LM = lean mass, FM = fat mass. For B&C, a non-parametric one-way ANOVA with multiple comparisons correction (Dunnet’s) was used.

**Table 2:** Echocardiography measurements in WT and KO male and female mice. Echocardiography measurements performed in the same animal at baseline (pre-procedure) and 8-weeks after undergoing sham or TAC surgery. Group numbers (n=9 for female, n=15 for male) were subject to whether the majority of data points were collected for a given animal at pre- and post-procedure (see methods for exclusion and inclusion criteria). Overall group size was smaller for females due to more severe disease being observed at 8-weeks post-TAC. # indicates that two mice were excluded due to heart rates being outside pre-specific exclusion criteria (450-650bpm). Bold text highlights values where there was a significant difference between WT and KO. Data are shown as mean±SEM. *p<0.05 vs WT TAC, †<0.05 vs sham of the same genotype, two-way ANOVA followed by Tukey post-hoc testing. Abbreviations: BW: Body Weight, (bpm): beats per minute, AWd: Anterior Wall diameter, LVEDD: Left Ventricular End-Diastolic Dimensions, PWd: Posterior Wall diameter, LVESD: Left Ventricular End-Systolic Dimensions, FS%: Fractional Shortening Percent; LV: left ventricle.

|  | Baseline |  | 8 Weeks Post-Procedure |  |  |  |
| --- | --- | --- | --- | --- | --- | --- |
|  |  |  | Sham |  | TAC |  |
| **Female** | WT (n = 9) | KO (n = 9) | WT (n = 3) | KO (n = 3) | WT (n = 6) | KO (n = 6) |
| BW (g) | 21.6 ± 1.1 | 21.2 ± 1.0 | 26.7 ± 1.8 | 24.5 ± 1.1 | 24.1 ± 0.9† | 22.2 ± 0.8† |
| Heart Rate (bpm) | 524 ± 19 | 499 ± 18 | 546 ± 8 | 525 ± 42 | 542 ± 17 | 559 ± 5 |
| AWd (mm) | 0.52 ± 0.03 | 0.54 ± 0.02 | 0.69 ± 0.07 | 0.68 ± 0.08 | 0.86 ± 0.02† | 0.82 ± 0.05† |
| LVEDD (mm) | 4.09 ± 0.10 | 4.15 ± 0.05 | 3.61 ± 0.21 | 3.97 ± 0.21 | 4.31 ± 0.25† | 4.84 ± 0.23† |
| PWd (mm) | 0.64 ± 0.02 | 0.58 ± 0.02 | 0.79 ± 0.09 | 0.72 ± 0.07 | 0.96 ± 0.05† | 0.93 ± 0.04† |
| LVESD (mm) | 2.91 ± 0.10 | 3.04 ± 0.06 | 2.51 ± 0.19 | 2.72 ± 0.19 | 3.51 ± 0.31† | 4.33 ± 0.21† |
| FS (%) | 29 ± 1.7 | 27 ± 0.9 | 31 ± 1.5 | 31 ± 4.5 | 19 ± 2.7† | **11 ± 1.1\*†** |
| LV Mass (mg) | 77.52 ± 5.94 | 79.59 ± 3.98 | 88.58 ± 5.51 | 97.20 ± 8.12 | 158.25 ± 9.70† | **198.04 ± 13.47\*†** |
|  |  |  | Sham |  | TAC |  |
| **Male** | WT (n = 15) | KO (n = 15) | WT (n = 5) | KO (n = 4) | WT (n = 8<sup>#</sup>) | KO (n = 11) |
| BW (g) | 22.0 ± 1.0 | 23.1 ± 0.7 | 28.8 ± 1.2 | 30.8 ± 0.9 | 29.4 ± 0.4 | 29.9 ± 0.6 |
| Heart Rate (bpm) | 536 ± 12 | 518 ± 18 | 545 ± 27 | 605 ± 34 | 602 ± 16 | 597 ± 12 |
| AWd (mm) | 0.53 ± 0.02 | 0.48 ± 0.01 | 0.61 ± 0.06 | 0.68 ± 0.03 | 0.87 ± 0.03† | 0.90 ± 0.02† |
| LVEDD (mm) | 4.06 ± 0.10 | 4.13 ± 0.10 | 4.25 ± 0.13 | 3.99 ± 0.05 | 4.84 ± 0.16† | 4.33 ± 0.15† |
| PWd (mm) | 0.60 ± 0.02 | 0.60 ± 0.01 | 0.70 ± 0.05 | 0.71 ± 0.05 | 0.97 ± 0.04† | 0.99 ± 0.03† |
| LVESD (mm) | 2.86 ± 0.10 | 2.94 ± 0.11 | 3.10 ± 0.16 | 2.59 ± 0.14 | 3.87 ± 0.16† | **3.25 ± 0.15\*†** |
| FS (%) | 30 ± 1.0 | 29 ± 1.4 | 27 ± 1.7 | 35 ± 3.7 | 20 ± 1.3† | **25 ± 1.1\*†** |
| LV Mass (mg) | 76.34 ± 3.51 | 75.64 ± 3.48 | 102.57 ± 15.00 | 97.14 ± 5.44 | 195.40 ± 11.73† | 172.54 ± 12.50† |

### Female OIP5-AS1 KO mice develop exacerbated disease in a pressure overload induced model of heart failure

Since there were no basal alterations in cardiac function in OIP5-AS1 KO mice, we sought to investigate whether inducing cardiac stress might reveal a phenotypic difference between the genotypes. Given that we observed reductions in expression of OIP5-AS1 in the setting of aortic constriction (AC) induced pressure overload in WT mouse hearts as shown in Figure 1G, we chose to subject OIP5-AS1 KO mice to a similar pressure overload procedure. Pressure overload also represents an appropriate model to test the effect of OIP5-AS1 deletion because it induces high-energy demand on the heart, thus directly impacting on the energetic pathways predicted to be associated with OIP5-AS1 function. For these studies, we performed transverse aortic constriction (TAC), which progressively induces heart failure over an approximately 8-week period. We performed TAC and sham surgery on both male and female weight matched OIP5-AS1 WT and KO mice at 7-10 weeks of age, and the phenotype was monitored over the ensuing 8 weeks (**Figure 3A**). Because the mice were weight matched, the aortic diameter and thus constriction at the time of surgery were assumed to be equivalent. Indeed, intra-cardiac and aortic catheter pressure analysis confirmed an equivalent pressure increase in a subset of WT and KO animals (**Supplemental Figure S2A**). Consistent with our previous studies [15,16,43,44], we demonstrated that the TAC procedure promoted significant pathological cardiac hypertrophy in WT mice following 8 weeks of TAC in both male and female mice compared to sham operated animals (**light grey bars, Figure 3B–3F**). Specifically, TAC operated WT mice (light grey bars) demonstrated an increase in total heart, left ventricle (LV), right ventricle (RV), atria and lung weights compared to sham operated animals (white bars), consistent with cardiomyopathy and congestive heart failure. With regards to OIP5-AS1 deletion, we demonstrated that female OIP5-AS1 KO mice (dark grey bars) had a more severe pathological hypertrophy and heart failure phenotype post-TAC than female WT mice, as indicated by significantly (p<0.05) increased heart weight, LV and RV weight, atrial weight and substantially heavier lung weight (**Figures 3B–3F**), the latter representing advanced cardiomyopathy and congestive heart failure. Interestingly, the same differences between genotypes were not observed in male mice, despite male WT mice displaying significant pathological cardiac hypertrophy (increased heart, LV and atria weights) following the TAC procedure.

**Figure 3:**
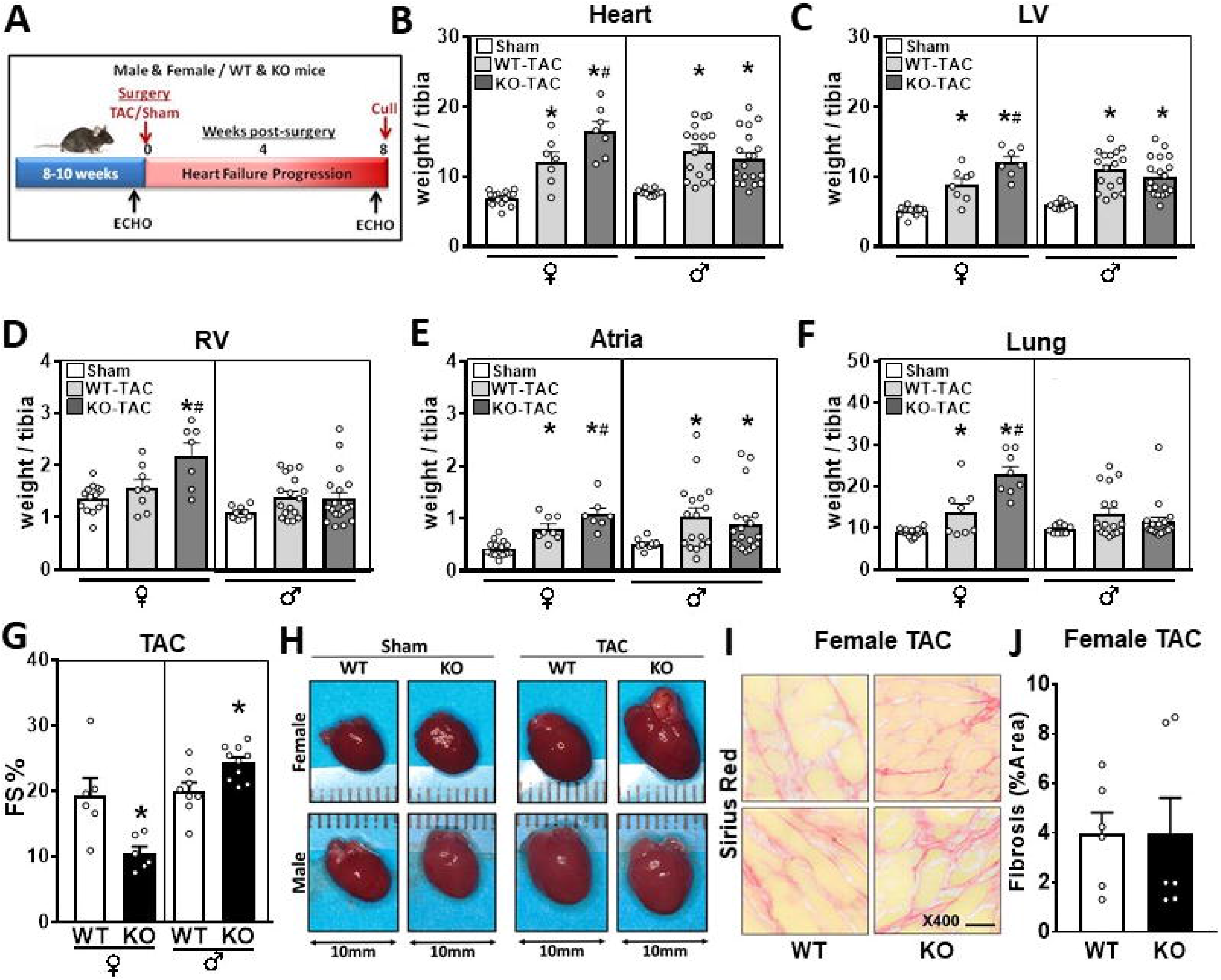
Female OIP5-AS1 mice demonstrated a worsened heart failure phenotype following TAC surgery. **A**. Schematic of experimental (TAC or Sham surgery) protocol performed on male and female WT and KO mice, which were followed for a further 8 weeks before cull. Tissue weights from cull were normalised to tibia length for **B**. whole heart weights, **C**. left ventricle **D**. right ventricle **E**. atria and **F**. whole lung (♀ = female, ♂ = male, female n=7-14/group, male n=9-19, mean±SEM, * p<0.05 versus sham, # p<0.05 versus WT-TAC of the same sex) **G**. Percent fractional shortening (FS %) as determined by echocardiography in male and female WT and KO mice at 8 weeks post-TAC procedure (n=6-11/group, mean±SEM, * p<0.05 versus sham for each sex). **H**. Representative photomicrographs of whole hearts from female and male WT and KO mice 8-weeks post procedure. **I**. Representative images of picrosirius red stained LV sections from female WT and KO mice post-TAC procedure, 400x magnification, scale bar = 20μM. **J**. Quantification of percent area picrosirius red staining (fibrosis) of entire LV section from WT and KO female mice undergoing TAC surgery (n=6/group). For panels B-G and panel J, a non-parametric one-way ANOVA with multiple comparisons correction (Dunnet’s) was used to test for significance, * p<0.05.

Echocardiography analysis demonstrated significant (p<0.05) alterations in LV wall thickness, dimensions and function in mice that had undergone TAC, compared to sham surgery mice, in both female and male animals (**Table 1**). Female OIP5-AS1 KO mice demonstrated a more pronounced decline in fractional shortening (FS%) following TAC compared to WT mice (11±1.1% vs 19±2.7%, respectively) (**Figure 3G**), indicating a worsened left ventricular function in female animals. Male OIP5-AS1 KO mice did not show a further decline in LV function in comparison to male WT mice post-TAC as demonstrated by FS% (**Table 2 and Figure 3G**). Photomicrographs of the hearts from male and female mice confirmed the increase in heart size in TAC-induced mice, and demonstrated the significant enlargement in both the ventricles and atria of KO mice, particularly in female mice (**top row, Figure 3H**). These pathological effects in the heart were associated with a greater incidence of clinical phenotypes of organ congestion, as indicated by an increase in lung weights in female KO mice in Figure 3F, and a greater percentage of female KO mice exhibiting thrombi in their atria compared to WT mice (55% vs 0%; shown visually in **Figure 3H**). Liver and kidney weights were equivalent between WT and KO male and female mice (**Supplemental Figure S2B&C**), whilst spleen weights were significantly larger in female KO TAC treated mice, potentially reflecting a general loss of health in these animals (**Supplemental Figure S2D**). We did not observe any alterations in fibrosis between WT and KO female mice as determined by picrosirius red staining of LV sections, which demonstrated no significant difference in percent area of fibrosis between WT and KO female mice following TAC surgery (**Figures 3I&3J**).

Collectively, these data demonstrated that female, but not male OIP5-AS1 KO mice displayed a more severe heart failure phenotype following pressure-induced cardiac-overload compared to wild type mice.

### Loss of OIP5-AS1 leads to transcriptional changes in gene networks associated with cardiomyopathy and mitochondrial function

In light of the exacerbated heart failure phenotype observed in female KO mice, we investigated transcriptional pathways that might be altered in OIP5-AS1 male and female mice. We did not observe differences in the basal expression of OIP5-AS1 in left ventricles between male and female WT mice, although the KO was equivalent between sexes (**Figure 4A**). We also observed no difference in OIP5-AS1 abundance between male and female WT mice following TAC (**Figure 4B**), as determined by RNA-seq. These results suggest that sex specific factors such as circulating hormones are most likely not influencing the expression of OIP5-AS1 in the hearts of WT mice either basally or after TAC, and thus more detailed analyses were necessary to tease out the potential causal pathways.

**Figure 4:**
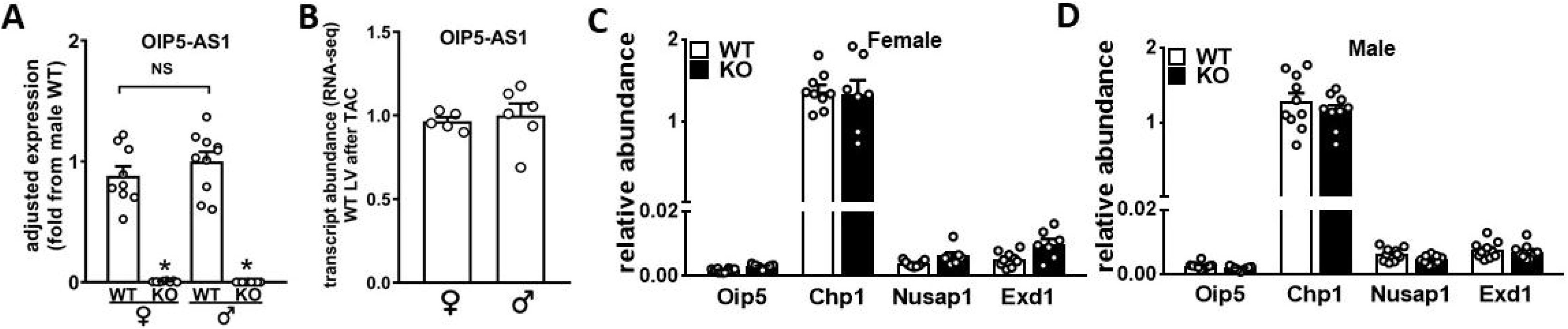
Expression of OIP5-AS1 and its neighbouring genes are not influenced by sex or *cis-* specific transcriptional mechanisms. **A**. Abundance of OIP5-AS1 transcripts as determined by qPCR in LV of male and female WT and OIP5-AS1 KO mice (n=9-10/group, mean±SEM, * p<0.05 versus WT, NS = not significant between male and female WT mice). **B**. Abundance of OIP5-AS1 transcripts as determined by RNA-seq in female and male WT LVs following TAC. Expression of OIP5-AS1 neighbouring genes (*Oip5, Chp1, Nusap1* and *Exd1*) in sham treated mice in **C**. female and **D**. male WT and OIP5-AS1 KO hearts as determined by qPCR (n=7-9/group, mean±SEM). Non-parametric one-way ANOVA with multiple comparisons correction (Dunnet’s) was used to test for significance, * p<0.05.

Some lncRNAs are proposed to function via a *cis* acting mechanism, whereby active transcription at the locus of a lncRNA, recruits transcriptional machinery and promotes opening of local chromatin and facilitating transcription of neighbouring genes. However, we did not observe any difference between WT and KO female or male mice in the expression of the four neighbouring genes of OIP5-AS1 (two proximal, two distal; *Oip5, Chp1, Nusap1* and *Exd1*) (**Figure 4C and 4D**), suggesting that the OIP5-AS1 phenotype is unlikely to be secondary to dysregulation of *cis* acting mechanisms. Thus, given that none of these archetypical pathways appeared to explain the mechanism by which OIP5-AS1 was functioning, we performed transcriptomic analysis on the WT and KO hearts in order to obtain a global overview of transcriptional differences between the genotypes.

For these analyses we performed RNA-sequencing on left ventricles (LV) from male and female WT and KO mice, all of which had undergone TAC surgery (n=6/group). Initial quality control analyses including PCA demonstrated segregation of the four groups (**Supplemental Figure S3A**). Consistent with *in vivo* phenotyping data, the greatest changes in differential gene expression were observed in female KO mice, whether it be compared to female WT mice (**Supplemental Figure S3B**) or Male KO mice (**Figure 5A)**. The number of genes significantly (q<0.1) altered between these groups was 66 genes between female WT and KO hearts, and ~1700 genes between female KO and male KO hearts (q<0.1) (**Figure 5A**). This latter finding may not seem si surprising, given that it is a comparison between male and female KO mice. However, when we compared the transcriptomes between male and female WT hearts, there were only 8 genes that were significantly (q<0.1) different, including known sex specific transcripts such as *Xist*, *Kdm5c* and *Ddx3x* (**Supplemental Figure S3C**). Thus, this indicated that the large transcriptional effect of OIP5-AS1 deletion between KO female and KO male hearts (~1700 genes) was not purely due to sex effects alone, but was a likely interaction between sex and the loss of OIP5-AS1 expression. Only one gene was significantly (q<0.1) altered between male WT and KO mice, and that was OIP5-AS1 itself (**Supplemental Figure S3D**). Thus, these data provide evidence that the loss of OIP5-AS1 results in specific alterations in cardiac gene expression post-TAC only in female mice.

**Figure 5:**
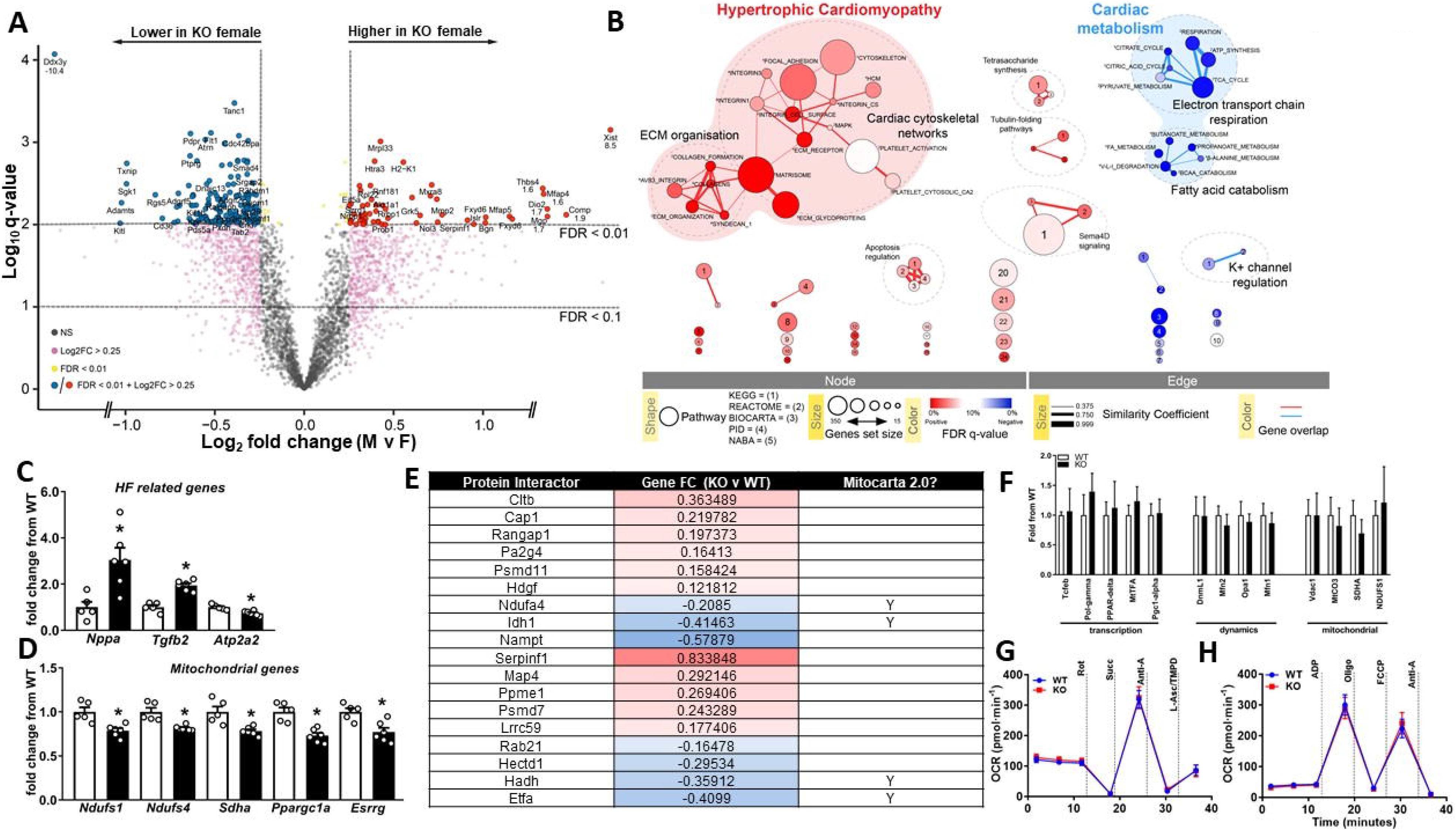
Transcriptomic and Integrative Analysis of OIP5-AS1 KO Hearts Reveals Alterations in Pathways Associated with Cardiomyopathy, Mitochondrial Metabolism and WNT Signalling. **A**. Volcano plot of LV gene expression analysed by RNA-seq that were differentially regulated between female (F) and male (M) KO mice following TAC (n=5-6/group), grey dots = non-significant, magenta = >0.25 log2FC, yellow dots = FDR<0.01, blue/red dots = >0.25log2FC and FDR<0.01. Off-scale points are specifically annotated with log2FC values at the individual data point. **B**. Functional map of genes altered in OIP5-AS1 KO female mice compared to WT, in LV following TAC. Colours indicate the association to either a positive (red = higher in KO) or negative (blue = lower in KO) enrichment. Node size is proportional to the total number of genes in each set. Edge thickness represents the similarity coefficient between gene/pathway sets (circle). Significance (FDR q-value as a percentage) of the enrichment is represented as a colour gradient, where a fuller color is more significant. The major functional groups are highlighted by the shaded background bubbles - enriched in KO (red); reduced in KO (blue). **C**. Abundance of transcripts in WT and OIP5-AS1 KO female mice as determined by RNA-sequencing in LV following TAC for genes related to heart failure (HF) (*Nppa, Tgfb2* and *Atp2a2*) and **D**. Related to mitochondrial function (*Ndufs1, Ndufs4, Sdha, Ppargc1a* and *Esrrg*), **E**. Table of proteins and genes that were shown to interact with OIP5-AS1 in a previously published dataset [46] and are also significant DEGs in our RNA-seq analysis. Targets were cross reference against MitoCarta 2.0 to identify those that are mitochondrial enriched (indicated by a “Y”). Fold change is represented with a gradient (blue = decreased in KO, red = increased in KO), where mitochondrial proteins show a strong trend to be decreased in OIP5-AS1 KO hearts. **F**. Gene expression of mitochondrial related genes as determined by qPCR analysis in left ventricles from female WT and KO sham operated mice (n=3/group). **G**. Oxygen consumption rate (OCR) as performed by Seahorse in mitochondria isolated from basal WT and OIP5-AS1 KO female hearts to test ETC Complex activity and to test **H**. Maximal respiratory capacity using the mitochondrial stress test (Rot = rotenone, Complex I inhibitor; Succ = Succinate, Complex II substrate; Anti-A = antimycin A, Complex III inhibitor; L-Asc/TMPD = L-ascorbate and Tetramethyl-p-phenylenediamine, Complex IV substrate; Oligo = Oligomycin, Complex V inhibitor; FCCP = Carbonyl cyanide 4-(trifluoromethoxy)phenylhydrazone, mitochondrial uncoupler) (n=7/group). For panels C and D, a non-parametric one-way ANOVA with multiple comparisons correction (Dunnet’s) was used to test for significance, * p<0.05.

To gain a better understanding of the pathways that were altered by OIP5-AS1 deletion, we performed pathway enrichment and network analysis on the gene sets altered between male and female KO mice. These studies demonstrated a notable enrichment (red networks) in genes associated with extracellular matrix (ECM)/cytoskeleton, and hypertrophic cardiomyopathy in female KO mice following TAC (**Figure 5B**). Moreover, we observed a substantial depletion (blue networks) in the expression of genes associated with cardiac metabolism in KO female hearts including electron transport chain (ETC) respiration, amino acid catabolism and fatty acid metabolism (**blue networks, Figure 5B**). Alterations in similar networks were also observed in the comparison of WT v KO female hearts, with changes in classic heart failure related genes such as *Nppa* (ANP), *Tgfb2* and *Atp2a2* (SERCA2) (**Figure 5C**). Moreover, changes in core components of complex I and II of the ETC were also observed including reductions in *Ndufs1*, *Ndufs4* and *Sdha* (**Figure 5D**). These findings were reassuring, because gene correlation analysis performed in Figure 1D suggested that OIP5-AS1 was likely associated with cardiomyopathy and mitochondrial/oxidative gene networks, predictions that appear to be supported by these analyses in KO mouse hearts. Moreover, our initial data in Figure 3I&J demonstrating that fibrosis pathways were not markedly different between WT and KO mice, was also supported by our RNA-seq data, with the majority of genes coding for collagens not substantially increased in female or male KO versus WT mice (**Supplemental Figure S3E and S3F)**.

Regarding specific mediators of this phenotype, it is interesting to note that two *bona fide* mitochondria related transcription factors were significantly decreased in OIP5-AS1 KO female hearts versus WT hearts. Specifically, we observed ~30% reduction in expression of *Ppargc1a* (PGC1alpha), a co-activator and regulator of genes important for mitochondrial biogenesis [22], and a ~25% reduction in *Esrrg* (ERRgamma) (**Figure 5D**). ERRgamma is a transcriptional co-activator that regulates transcription of genes important for oxidative capacity and mitochondrial function [45], and has also been shown to influence neonatal to adult transition in cardiomyocytes [37], an important time in development where we observed substantial increases in OIP5-AS1 expression (Figure 1).

These observed alterations in mitochondrial networks were of potential mechanistic interest. However, because we have no direct evidence for how OIP5-AS1 functions in cardiomyocytes, we instead mined previously published datasets in an attempt to gain further mechanistic insights. One such dataset was a protein interaction dataset, which used pull down approaches to identify proteins that interacted with OIP5-AS1 in ES cells [46]. By comparing their list of OIP5-AS1 interacting proteins with our differentially expressed gene set, we identified 18 genes/proteins that were consistent between the two datasets (**Figure 5E** and **Supplementary Table 3**). There were a mix of genes upregulated (red shading) and downregulated (blue shading) in the KO heart, however it was apparent that the majority of the down regulated genes (blue shading) were known mitochondrial associated proteins as indicated by their presence in the MitoCarta 2.0 database[47] (indicated by a “Y” in the table). Thus, we hypothesise that OIP5-AS1 may interact with critical proteins from the mitochondria (e.g. Ndufa4, Idh1, Hadh, Etfa), and that loss of OIP5-AS1 leads to a disruption of this function. This effect is likely to be stress specific, such as that induced by TAC, because we also demonstrate that the expression of these mitochondrial genes were not different between WT and KO female hearts in the basal (i.e. sham) setting (**Figure 5F**), nor was mitochondrial function different in the basal setting as measured by Seahorse (**Figure 5G&H**). Thus, OIP5-AS1 loss potentially alters mitochondrial networks in the setting of stress in the female heart, yet loss of OIP5-AS1 in a basal, unstressed setting does not impact mitochondrial function.

Collectively, our findings provide evidence that disruption OIP5-AS1 leads to an exacerbated heart failure progression following stress in female mice, a phenotype that may be linked to specific components of the mitochondrial network.

## Discussion

In the current study we have described and characterised the lncRNA OIP5-AS1 to be enriched and functional in cardiac muscle. OIP5-AS1 demonstrates several regions of high conservation between mouse and human in its nucleotide sequence, and thus its function and roles are likely to be conserved. Upon generating and studying a novel OIP5-AS1 KO mouse model, we demonstrated that loss of OIP5-AS1 renders female mice more prone to cardiac overload induced heart failure, a phenotype not observed in male mice. Transcriptomic and integrative analyse indicated that OIP5-AS1 modulates molecular regulators of mitochondrial function, exemplified by alterations in key mitochondrial gene sets both in cardiomyocytes isolated from mouse hearts, and in OIP5-AS1 KO heart tissue. Interestingly, we did not observe alterations in mitochondrial gene expression or mitochondrial function in non-stressed hearts, suggesting that the changes observed in stressed KO hearts were a maladaptive response. These data are consistent with the phenotype being primarily driven by changes in metabolism of cardiomyocytes in female OIP5-AS1 KO mice, rather than extracellular matrix remodelling such as scarring and fibrosis. The phenotype is also reminiscent of congenital defects of mitochondrial dysfunction that often precipitates as cardiomyopathy and heart failure in the absence of significant fibrosis.

Our transcriptional analyses identified alterations in two mitochondrial transcription factors (ERRgamma and PGC1alpha) in the female TAC KO heart. This may represent one mechanism by which OIP5-AS1 KO mice exhibit a worsening heart failure phenotype in a setting of pressure overload. ERRgamma was specifically down-regulated in female OIP5-AS1 KO hearts following chronic cardiac pressure overload, a pathological setting that increases energy demand and thus aids in revealing a contractile dysfunction phenotype. ERRgamma is a transcriptional regulator of genes important for energy production including mitochondrial activity [45]. Evidently, others have demonstrated that ERRgamma is an essential co-ordinator of cardiac metabolism and function [48].

Consistent with an effect on mitochondrial dysfunction, we also observed a significant reduction in PGC1alpha expression, a critical regulator of mitochondrial biogenesis pathways [49]. Given the significant loss of both PGC1alpha and ERRgamma, it is likely that the hearts of female OIP5-AS1 KO mice would have an impaired ability to generate energy and thus deprive the failing heart of ATP, driving a more severe phenotype in the absence of fibrosis. Indeed, this is important given that other studies have proposed that a 30-40% loss of PGC1alpha is sufficient to drive exacerbated heart failure [48,50,51]. Moreover, similar to our findings, studies by Chang *et al* [51] and Warren *et al* [52] demonstrated that dystrophic and Smyd1 driven cardiomyopathy respectively, were characterized by a dysregulation of mitochondrial biogenesis and function, exemplified by a downregulation of PGC1alpha.

Previous studies have demonstrated that lncRNAs can directly influence the expression of nuclear and mitochondrial encoded genes, with loss of lncRNAs leading to reduced mitochondrial activity and pathological phenotypes [53–55]. Thus, OIP5-AS1 may interact with transcriptional complexes harbouring PGC1alpha or ERRgamma that directly regulate the expression of these genes. Our findings are consistent and supportive of the work recently published by Niu and colleagues who demonstrated that overexpression of OIP5-AS1 in rat hearts and cardiac cell cultures was protective against ischemia reperfusion injury resulting from transient myocardial infarction [38]. They proposed that this protection was mediated by changes in mitochondrial function driven in part by alterations in the PGC1alpha axis. Thus these findings, combined with our new data generated in a novel KO mouse model, provide strong evidence for OIP5-AS1 regulating a conserved cardiac energetics program in the heart.

Regarding the specific mechanisms of OIP5-AS1 function, work by Kleaveland *et al* [28] demonstrated that OIP5-AS1 in the brain was predominantly localised to the cytoplasm (73%), where OIP5-AS1 (Cyrano) acted as a sponge for miRNA-7, subsequently decreasing the nuclear abundance of a circular RNA called *Cdr1-AS*. Others have demonstrated that OIP5-AS1 interacts with the competing endogenous RNA (ceRNA), HuR (*Elavl1*) in HeLa cells, alters the abundance of HuR (*Elavl1*) complex [29], or binds and effects the activity of miR-29a[38]. Whether these mechanisms are active in our model in the mouse heart is not known, however our RNA-seq analysis demonstrates that neither miRNA-7, miR-29a or *Elavl1* were altered in hearts of female KO mice. These discrepancies might be explained by divergent actions of OIP5-AS1 in striated muscle compared to other tissues, or that our transcriptome data was not optimised to capture miRs efficiently. It may also be related to the method in which OIP5-AS1 was deleted/silenced, or that our KO studies were performed in a pathological setting and the majority of other studies were not.

Regarding the mouse models, Kleaveland *et al* generated mice that deleted a specific region of exon 3 of Cyrano (OIP5-AS1) through cre-lox recombination, whereas we used CRISPR/Cas9 to delete the entire gene locus. Kleaveland *et al* also demonstrated mild subclinical *cis*-effects in the brain of their KO model, namely a small but significant increase in the expression of the neighbouring gene *Nusap1*, whereas we did not observe any change in *Nusap1* expression, at least not in the heart. Each of these models comes with their own limitations. For example, in our model there is the possibility we have removed other non-coding or coding transcripts present in the region, however given that no other genes or ncRNAs are annotated to the locus - this is unlikely. We may also have inadvertently removed important sites that are necessary for chromatin looping, or other long range chromatin interactions[56], however again, without having specific knowledge of this it cannot be tested directly. With regard to the model by Kleaveland et al, many regions of Cyrano have been shown to be important for its function. Specifically, Smith et al demonstrated that exon 3 of OIP5-AS1 is important for protein binding at multiple sites, and therefore removing only a small fraction of exon 3 is likely to impact very specific OIP5-AS1 mechanisms (which was likely the desired goal), which may not induce an overt phenotype. Similarly, the in vivo data presented by Niu et al in the setting of MI in rats remains somewhat inconclusive, because the authors use AAVs to overexpress OIP5-AS1, which have a packaging limit of ~5-6kb. Given that the major transcript of OIP5-AS1 (accounting for >90% of total OIP5-AS1 in the heart) is well above this limit at ~9.5kb, this suggests that a truncated version of OIP5-AS1 was overexpressed, raising questions about the interpretation and translatability of their findings.

Nevertheless, with each model there remains specific advantages and disadvantages and thus each of these approaches contribute unique knowledge to the field. An alternative approach to studying lncRNAs has been to insert loxP sites that flank the promoter region of the lncRNA, which would remove its ability to be transcriptionally regulated; however, no studies have yet been published using such an approach to study OIP5-AS1.

Another important distinction between our study and those from other groups is that we investigated both sexes, and observed a phenotype only in female mice. We also studied these animals in a chronic disease setting that directly increases load on the heart, similar to that which occurs with hypertension and other cardiovascular conditions, which may be required to reveal specific phenotypes. Although not explicitly stated, it is assumed that previous groups only studied male OIP5-AS1 KO mice, possibly explaining in part the discrepancies observed in molecular phenotypes between our KO study and others.

In conclusion, the major findings from this study include: 1) use of experimental and bioinformatic analyses of cardiac tissue and cells from both murine and human origin to confirm OIP5-AS1 as cardiac muscle enriched lncRNA, 2) generation and characterisation of a novel mouse model with deletion of OIP5-AS1, 3) uncovering a critical role of OIP5-AS1 in the female heart in a setting of cardiac pressure overload and 4) in-depth transcriptional analyses highlighting a dysregulation of mitochondrial genes in the female heart in the setting of pressure overload.

Future studies would look to gain further mechanistic insight, as it will be important to directly assess cardiac metabolism in KO mice, and ascertain the contribution of a mitochondrial defect in contributing to the accelerated heart failure phenotype in female TAC KO mice. In addition, given our data demonstrating that OIP5-AS1 is significantly reduced in the setting of cardiac disease, it will be interesting to determine whether overexpression or reconstitution of OIP5-AS1 in female mice is effective at protecting against heart failure, as was demonstrated in rat models of MI. Finally, as OIP5-AS1 is also enriched in skeletal muscle and the brain, studies investigating these tissues in our KO model would also be of interest.

In summary, our study sheds light on our current understanding of sexual dimorphism observed in heart disease by demonstrating the female-sex restricted regulation of cardiac maladaptation by lncRNA OIP5-AS1. This is of particular importance considering the clinical observation that women are up to 4-fold more likely to develop heart failure in some settings than men, with a poor understanding of the molecular underpinnings of this observation. Our data suggests that future studies into sex disparities in heart failure progression in humans, might consider analysing mitochondrial function and lncRNAs such as OIP5-AS1 as being potential regulators.

## Acknowledgements

We acknowledge funding support from the Victorian State Government OIS program to Baker Heart & Diabetes Institute (BHDI). These studies were supported in part by funding from the National Health & Medical Research Council (NHMRC) of Australia to BD and JM (APP1127336). BD and AC received funding from the National Heart Foundation of Australia, via the Future Leader Fellowship Scheme (101789 and 100067, respectively). JM and PG are Research Fellows of the NHMRC (APP1078985 and APP1117835, respectively). We acknowledge the use of facilities and technical assistance from the Monash Histology Platform, Department of Anatomy and Developmental Biology, Monash University, and use of the gene set enrichment analysis (GSEA) software, and Molecular Signature Database (MSigDB) [23]. We thank all members of the Cardiac Hypertrophy, MMA and LMCD laboratories at BHDI for their ongoing contributions.

## Author Contributions

BD & JM conceived the study and wrote the manuscript. BD, JM, AC and XJD designed the mouse studies and directed experimental analysis. BD, JM, AC, AZ, SL, YL, SB, SM, EG, and SRC analysed data and interpreted findings. BD, SL, JM, HK, YL, AZ, DD, XMG and XJD performed animal experiments and analysed data. BD, JM, AC, TC, TV, ET, JH, GQR, EP, PG and MI provided resources, data analysis, experimental guidance and reagents. All authors had the opportunity to read, edit and identify points for clarification before submission.

---

**Supplementary Figure S1:**
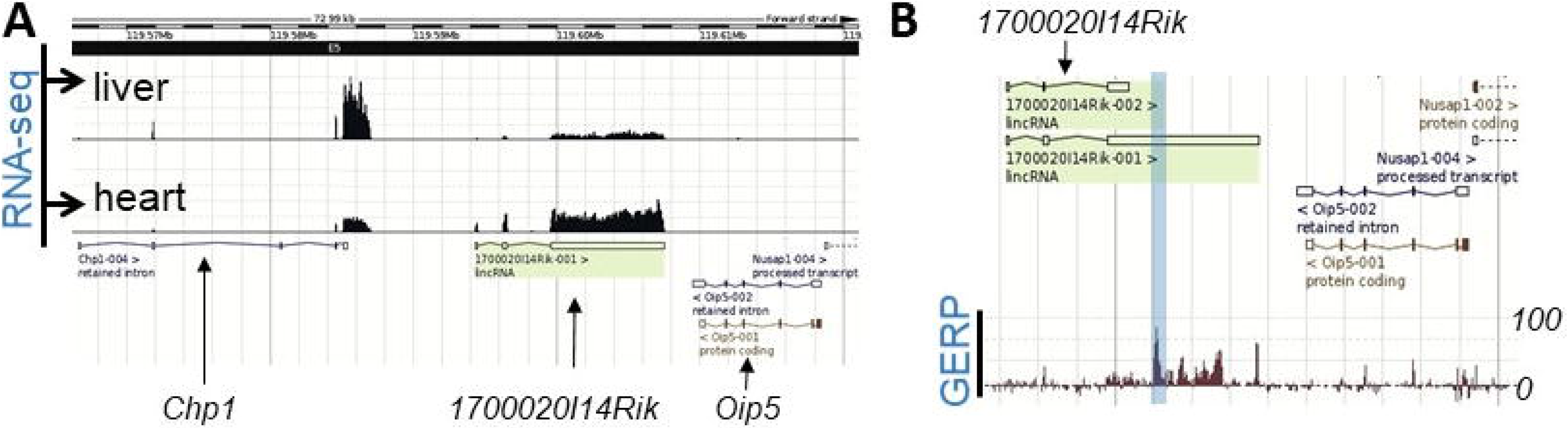
Location, expression and homology of OIP5-AS1. **A**. *Ensembl* sourced RNA-sequencing tracks of transcript expression in mouse liver and heart at the OIP5-AS1 locus. **B**. *Ensembl* sourced Genomic Evolutionary Rate Profiling (GERP) data across the OIP5-AS1 gene locus. A greater amplitude in the GERP peak indicates a greater conservation of that region across vertebrate species. Blue box indicates the region of 100% homology in exon 3.

**Supplementary Figure S2:**
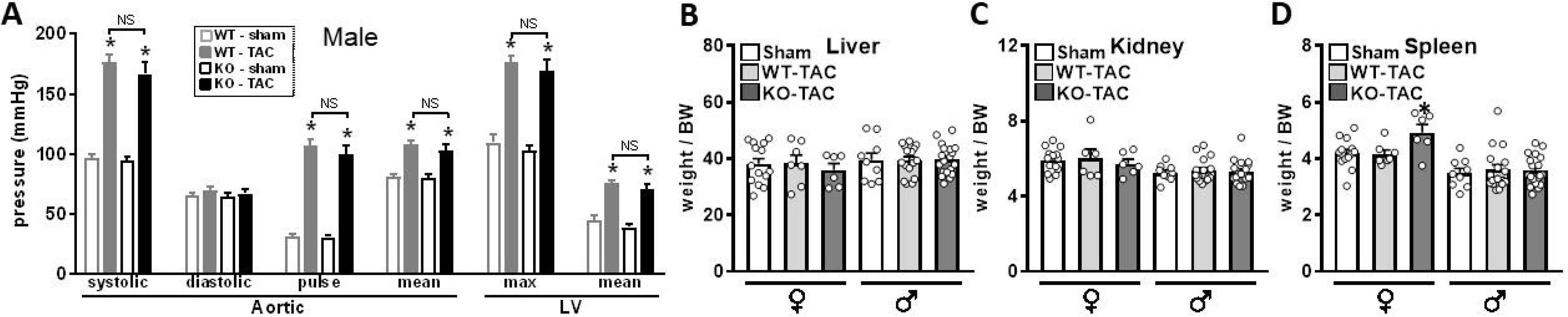
Additional phenotyping data from WT and KO mice undergoing Sham and TAC surgery. **A**. Hemodynamic catheter measurements recorded from the aorta and left ventricle (LV) of sham and TAC operated WT and OIP5-AS1 KO male mice (n=3-11/group, mean±SEM), * p<0.05 versus genotype equivalent sham, NS = not significant between WT and KO TAC treated animals; Organ weights adjusted to body weight (BW) in male and female mice 8-weeks post procedure for **B**. liver, **C**. kidney and **D**. spleen, (n=7-13/group, mean±SEM, * p<0.05 versus sham for relevant sex). Weights for these organs were adjusted to BW.

**Supplementary Figure S3:**
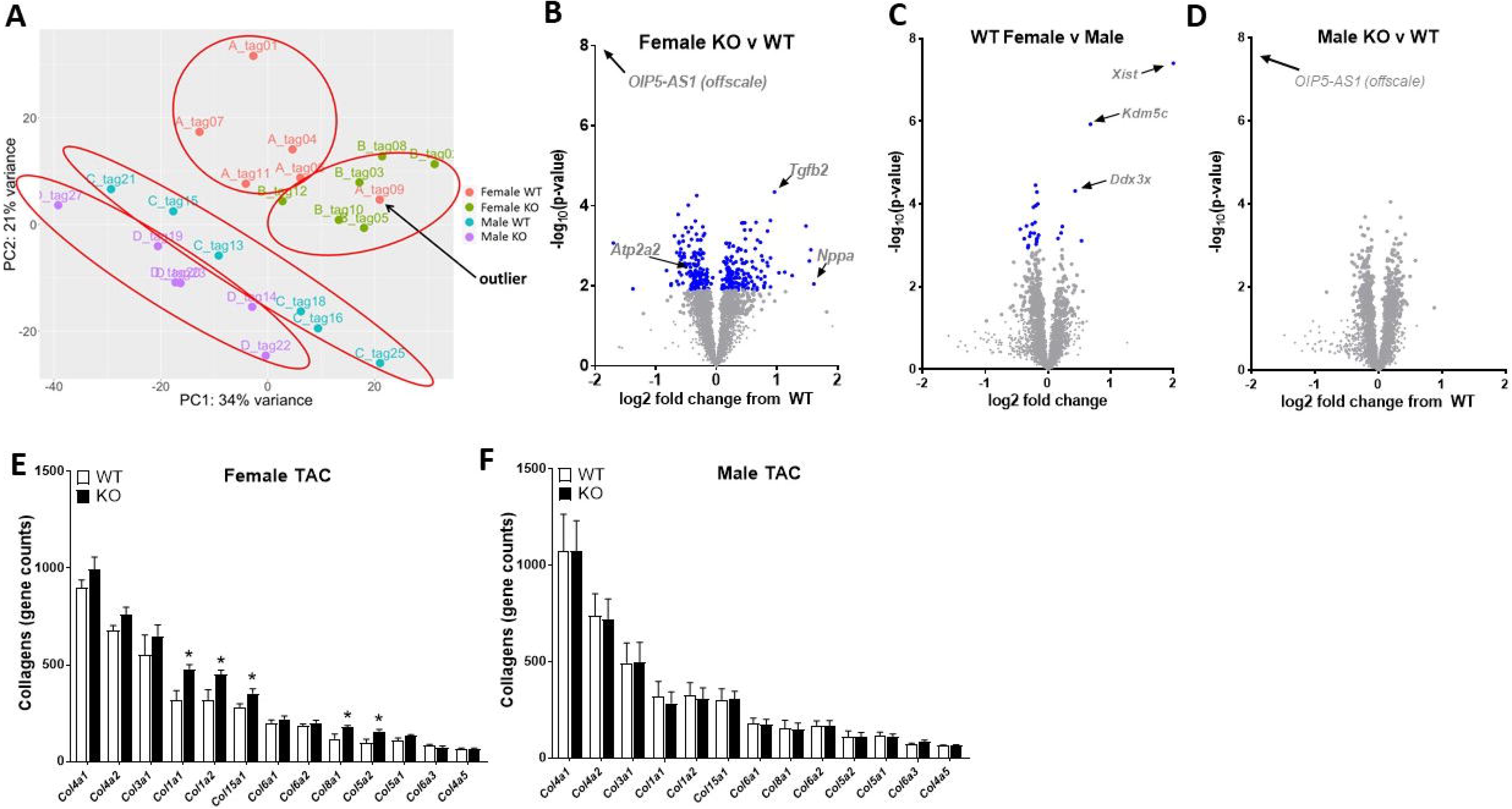
Accompanying RNA-sequencing Data Analysis. **A**. Principle component analysis (PCA) of RNA-sequencing data demonstrating separation of the four groups of samples based on variance (red circles). One sample from the “Female WT” group demonstrated an inconsistent variance compared to its other group members indicating a technical abnormality (black arrow – “outlier”) and was subsequently removed from enrichment and network analysis. **B**. Volcano plot of genes regulated in LV following TAC between female KO and WT mice (n=5/group for female WT, n=6/group for female KO), grey dots = not-significant, blue dots = FDR<0.1). Arrows demonstrate genes known to be regulated in the setting of heart failure. **C**. Volcano plot of genes regulated in LV following TAC between female and male WT mice (n=5/group for female WT, n=6/group for male WT), grey dots = not-significant, blue dots = FDR<0.1). Arrows demonstrate known sexual dimorphic genes. **D**. Volcano plot of genes regulated in LV following TAC between male WT and KO mice (n=6/group), grey dots = not-significant. Arrow indicates that change in OIP5-AS1 expression is not shown as it is off scale. Absolute transcript abundance of all collagen related genes in LV from RNA-seq data in **E**. female WT and KO mice following TAC and **F**. male WT and KO mice following TAC (* p<0.05 versus WT, n=5-6/group).

## References

1. Cech TR, Steitz JA (2014) The noncoding RNA revolution-trashing old rules to forge new ones. Cell 157: 77–94.

2. Kashi K, Henderson L, Bonetti A, Carninci P (2016) Discovery and functional analysis of lncRNAs: Methodologies to investigate an uncharacterized transcriptome. Biochim Biophys Acta 1859: 3–15.

3. Mallory AC, Shkumatava A (2015) LncRNAs in vertebrates: advances and challenges. Biochimie 117: 3–14.

4. Ulitsky I, Bartel DP (2013) lincRNAs: genomics, evolution, and mechanisms. Cell 154: 26–46.

5. St Laurent G, Wahlestedt C, Kapranov P (2015) The Landscape of long noncoding RNA classification. Trends Genet 31: 239–251.

6. McMullen JR, Drew BG (2016) Long non-coding RNAs (lncRNAs) in skeletal and cardiac muscle: potential therapeutic and diagnostic targets? Clin Sci (Lond) 130: 2245–2256.

7. Klattenhoff CA, Scheuermann JC, Surface LE, Bradley RK, Fields PA, et al. (2013) Braveheart, a long noncoding RNA required for cardiovascular lineage commitment. Cell 152: 570–583.

8. Colley SM, Leedman PJ (2011) Steroid Receptor RNA Activator - A nuclear receptor coregulator with multiple partners: Insights and challenges. Biochimie 93: 1966–1972.

9. Grote P, Wittler L, Hendrix D, Koch F, Wahrisch S, et al. (2013) The tissue-specific lncRNA Fendrr is an essential regulator of heart and body wall development in the mouse. Dev Cell 24: 206–214.

10. Ounzain S, Micheletti R, Beckmann T, Schroen B, Alexanian M, et al. (2015) Genome-wide profiling of the cardiac transcriptome after myocardial infarction identifies novel heart-specific long non-coding RNAs. Eur Heart J 36: 353–368a.

11. Han P, Li W, Lin CH, Yang J, Shang C, et al. (2014) A long noncoding RNA protects the heart from pathological hypertrophy. Nature 514: 102–106.

12. Ishii N, Ozaki K, Sato H, Mizuno H, Saito S, et al. (2006) Identification of a novel non-coding RNA, MIAT, that confers risk of myocardial infarction. J Hum Genet 51: 1087–1099.

13. Kumarswamy R, Bauters C, Volkmann I, Maury F, Fetisch J, et al. (2014) Circulating long noncoding RNA, LIPCAR, predicts survival in patients with heart failure. Circ Res 114: 1569–1575.

14. Wang K, Liu F, Zhou LY, Long B, Yuan SM, et al. (2014) The long noncoding RNA CHRF regulates cardiac hypertrophy by targeting miR-489. Circ Res 114: 1377–1388.

15. Gao XM, Kiriazis H, Moore XL, Feng XH, Sheppard K, et al. (2005) Regression of pressure overload-induced left ventricular hypertrophy in mice. Am J Physiol Heart Circ Physiol 288: H2702–2707.

16. Kiriazis H, Wang K, Xu Q, Gao XM, Ming Z, et al. (2008) Knockout of beta(1)- and beta(2)-adrenoceptors attenuates pressure overload-induced cardiac hypertrophy and fibrosis. Br J Pharmacol 153: 684–692.

17. Du XJ, Autelitano DJ, Dilley RJ, Wang B, Dart AM, et al. (2000) beta(2)-adrenergic receptor overexpression exacerbates development of heart failure after aortic stenosis. Circulation 101: 71–77.

18. Donner DG, Kiriazis H, Du XJ, Marwick TH, McMullen JR (2018) Improving the quality of preclinical research echocardiography: observations, training, and guidelines for measurement. Am J Physiol Heart Circ Physiol 315: H58–H70.

19. Bond ST, Kim J, Calkin AC, Drew BG (2019) The Antioxidant Moiety of MitoQ Imparts Minimal Metabolic Effects in Adipose Tissue of High Fat Fed Mice. Front Physiol 10: 543.

20. Bond ST, Moody SC, Liu Y, Civelek M, Villanueva CJ, et al. (2019) The E3 ligase MARCH5 is a PPARgamma target gene that regulates mitochondria and metabolism in adipocytes. Am J Physiol Endocrinol Metab 316: E293–E304.

21. Huang da W, Sherman BT, Lempicki RA (2009) Systematic and integrative analysis of large gene lists using DAVID bioinformatics resources. Nat Protoc 4: 44–57.

22. Mootha VK, Lindgren CM, Eriksson KF, Subramanian A, Sihag S, et al. (2003) PGC-1alpha-responsive genes involved in oxidative phosphorylation are coordinately downregulated in human diabetes. Nat Genet 34: 267–273.

23. Subramanian A, Tamayo P, Mootha VK, Mukherjee S, Ebert BL, et al. (2005) Gene set enrichment analysis: a knowledge-based approach for interpreting genome-wide expression profiles. Proc Natl Acad Sci U S A 102: 15545–15550.

24. Raudvere U, Kolberg L, Kuzmin I, Arak T, Adler P, et al. (2019) g:Profiler: a web server for functional enrichment analysis and conversions of gene lists (2019 update). Nucleic Acids Res 47: W191–W198.

25. Reimand J, Kull M, Peterson H, Hansen J, Vilo J (2007) g:Profiler--a web-based toolset for functional profiling of gene lists from large-scale experiments. Nucleic Acids Res 35: W193–200.

26. Schindelin J, Arganda-Carreras I, Frise E, Kaynig V, Longair M, et al. (2012) Fiji: an open-source platform for biological-image analysis. Nat Methods 9: 676–682.

27. Ulitsky I, Shkumatava A, Jan CH, Sive H, Bartel DP (2011) Conserved function of lincRNAs in vertebrate embryonic development despite rapid sequence evolution. Cell 147: 1537–1550.

28. Kleaveland B, Shi CY, Stefano J, Bartel DP (2018) A Network of Noncoding Regulatory RNAs Acts in the Mammalian Brain. Cell 174: 350–362 e317.

29. Kim J, Abdelmohsen K, Yang X, De S, Grammatikakis I, et al. (2016) LncRNA OIP5-AS1/cyrano sponges RNA-binding protein HuR. Nucleic Acids Res 44: 2378–2392.

30. Smith KN, Starmer J, Miller SC, Sethupathy P, Magnuson T (2017) Long Noncoding RNA Moderates MicroRNA Activity to Maintain Self-Renewal in Embryonic Stem Cells. Stem Cell Reports 9: 108–121.

31. Zheng D, Wang B, Zhu X, Hu J, Sun J, et al. (2019) LncRNA OIP5-AS1 inhibits osteoblast differentiation of valve interstitial cells via miR-137/TWIST11 axis. Biochem Biophys Res Commun 511: 826–832.

32. Diederichs S (2014) The four dimensions of noncoding RNA conservation. Trends Genet 30: 121–123.

33. Thorrez L, Van Deun K, Tranchevent LC, Van Lommel L, Engelen K, et al. (2008) Using ribosomal protein genes as reference: a tale of caution. PLoS One 3: e1854.

34. Ribas V, Drew BG, Zhou Z, Phun J, Kalajian NY, et al. (2016) Skeletal muscle action of estrogen receptor alpha is critical for the maintenance of mitochondrial function and metabolic homeostasis in females. Sci Transl Med 8: 334ra354.

35. Dey BK, Pfeifer K, Dutta A (2014) The H19 long noncoding RNA gives rise to microRNAs miR-675-3p and miR-675-5p to promote skeletal muscle differentiation and regeneration. Genes Dev 28: 491–501.

36. Watts R, Johnsen VL, Shearer J, Hittel DS (2013) Myostatin-induced inhibition of the long noncoding RNA Malat1 is associated with decreased myogenesis. Am J Physiol Cell Physiol 304: C995–1001.

37. Quaife-Ryan GA, Sim CB, Ziemann M, Kaspi A, Rafehi H, et al. (2017) Multicellular Transcriptional Analysis of Mammalian Heart Regeneration. Circulation 136: 1123–1139.

38. Niu X, Pu S, Ling C, Xu J, Wang J, et al. (2020) lncRNA Oip5-as1 attenuates myocardial ischaemia/reperfusion injury by sponging miR-29a to activate the SIRT1/AMPK/PGC1alpha pathway. Cell Prolif 53: e12818.

39. Kennedy TL, Swiderski K, Murphy KT, Gehrig SM, Curl CL, et al. (2016) BGP-15 Improves Aspects of the Dystrophic Pathology in mdx and dko Mice with Differing Efficacies in Heart and Skeletal Muscle. Am J Pathol 186: 3246–3260.

40. Chen JL, Walton KL, Hagg A, Colgan TD, Johnson K, et al. (2017) Specific targeting of TGF-beta family ligands demonstrates distinct roles in the regulation of muscle mass in health and disease. Proc Natl Acad Sci U S A 114: E5266–E5275.

41. McMullen JR, Amirahmadi F, Woodcock EA, Schinke-Braun M, Bouwman RD, et al. (2007) Protective effects of exercise and phosphoinositide 3-kinase(p110alpha) signaling in dilated and hypertrophic cardiomyopathy. Proc Natl Acad Sci U S A 104: 612–617.

42. McMullen JR, Sherwood MC, Tarnavski O, Zhang L, Dorfman AL, et al. (2004) Inhibition of mTOR signaling with rapamycin regresses established cardiac hypertrophy induced by pressure overload. Circulation 109: 3050–3055.

43. Bernardo BC, Gao XM, Winbanks CE, Boey EJ, Tham YK, et al. (2012) Therapeutic inhibition of the miR-34 family attenuates pathological cardiac remodeling and improves heart function. Proc Natl Acad Sci U S A 109: 17615–17620.

44. Bernardo BC, Nguyen SS, Winbanks CE, Gao XM, Boey EJ, et al. (2014) Therapeutic silencing of miR-652 restores heart function and attenuates adverse remodeling in a setting of established pathological hypertrophy. FASEB J 28: 5097–5110.

45. Alaynick WA, Kondo RP, Xie W, He W, Dufour CR, et al. (2007) ERRgamma directs and maintains the transition to oxidative metabolism in the postnatal heart. Cell Metab 6: 13–24.

46. Smith KN, Starmer J, Magnuson T (2018) Interactome determination of a Long Noncoding RNA implicated in Embryonic Stem Cell Self-Renewal. Sci Rep 8: 17568.

47. Calvo SE, Clauser KR, Mootha VK (2016) MitoCarta2.0: an updated inventory of mammalian mitochondrial proteins. Nucleic Acids Res 44: D1251–1257.

48. Wang T, McDonald C, Petrenko NB, Leblanc M, Wang T, et al. (2015) Estrogen-related receptor alpha (ERRalpha) and ERRgamma are essential coordinators of cardiac metabolism and function. Mol Cell Biol 35: 1281–1298.

49. Karkkainen O, Tuomainen T, Mutikainen M, Lehtonen M, Ruas JL, et al. (2019) Heart specific PGC-1alpha deletion identifies metabolome of cardiac restricted metabolic heart failure. Cardiovasc Res 115: 107–118.

50. Jia Y, Chang HC, Schipma MJ, Liu J, Shete V, et al. (2016) Cardiomyocyte-Specific Ablation of Med1 Subunit of the Mediator Complex Causes Lethal Dilated Cardiomyopathy in Mice. PLoS One 11: e0160755.

51. Chang AC, Ong SG, LaGory EL, Kraft PE, Giaccia AJ, et al. (2016) Telomere shortening and metabolic compromise underlie dystrophic cardiomyopathy. Proc Natl Acad Sci U S A 113: 13120–13125.

52. Warren JS, Tracy CM, Miller MR, Makaju A, Szulik MW, et al. (2018) Histone methyltransferase Smyd1 regulates mitochondrial energetics in the heart. Proc Natl Acad Sci U S A 115: E7871–E7880.

53. Vendramin R, Verheyden Y, Ishikawa H, Goedert L, Nicolas E, et al. (2018) SAMMSON fosters cancer cell fitness by concertedly enhancing mitochondrial and cytosolic translation. Nat Struct Mol Biol 25: 1035–1046.

54. Noh JH, Kim KM, Abdelmohsen K, Yoon JH, Panda AC, et al. (2016) HuR and GRSF1 modulate the nuclear export and mitochondrial localization of the lncRNA RMRP. Genes Dev 30: 1224–1239.

55. Long J, Badal SS, Ye Z, Wang Y, Ayanga BA, et al. (2016) Long noncoding RNA Tug1 regulates mitochondrial bioenergetics in diabetic nephropathy. J Clin Invest 126: 4205–4218.

56. Mumbach MR, Satpathy AT, Boyle EA, Dai C, Gowen BG, et al. (2017) Enhancer connectome in primary human cells identifies target genes of disease-associated DNA elements. Nat Genet 49: 1602–1612.

